# Regulation of sensorimotor serial learning in speech production by motor compensation rather than sensory error

**DOI:** 10.1101/2024.10.27.620480

**Authors:** Yuhan Lu, Xiaowei Tang, Zhenyan Xiao, Anqi Xu, Junxi Chen, Xing Tian

## Abstract

Motor control is essential for organisms to efficiently interact with the environment by maintaining accurate action and adjusting to future changes. Speech production, one of the most complex motor behaviors, relies on a feedback control process to detect sensory errors and trigger updates in a feedforward control process that implements compensations. However, the specific contributions of these critical processes in sensorimotor learning during continuous vocal production remain debated. Here, we used two experimental designs in five experiments to dissociate these mechanisms.

First, we employed a serial-dependence design with randomized pitch perturbations, dissociating the influences of sensory errors and motor compensation on subsequent vocalizations on a trial-by-trial basis. We found that motor compensation, rather than sensory errors, predicted the compensatory responses in the subsequent trials, suggesting instantaneous serial learning mediated by updates in the feedforward process. This compensation-driven serial learning was generalized across productions of different vowel categories. Second, we further implemented a serial-dependence adaptation design in a sentential context, where auditory perturbation occurred only on a preceding syllable. Any learning effects in its subsequent syllable without pitch perturbation would reflect changes in the speech motor representation. Our results consistently revealed that compensation in the preceding syllable predicted pitch changes in the subsequent syllable, but only when the two adjacent syllables were embedded within a word boundary. Collectively, the study provides ecological-valid evidence supporting that error-based motor compensation, incorporating cognitive and linguistic constraints, directly regulates the speech motor representation and mediates the instantaneous serial learning in successive actions.

## Introduction

The accuracy and precision of motor output, as a result of efficient interplay among motor, sensory and cognitive systems, are crucial for human survival and social interactions.^1-3^ In the domain of speaking, for example, the control of articulators relies on real-time integration among the language system, speech motor system and sensory feedback from the external environment.^4-7^ Specifically, the brain anticipates the sensory consequences of articulatory movements via internal forward models,^8-13^ continuously compares the predictions to sensory feedback in the feedback process (**Fig. 1A**, left panel).^14-16^ Experimentally introduced auditory perturbations (e.g., pitch shifts) create mismatches between expected and heard feedback, effectively serving as sensory errors that trigger the inverse model and update speech motor representation in the feedforward process to correct articulation in real-time (**Fig. 1A**, left panel).^17,18^ Notably, such online compensatory adjustments can propagate to influence subsequent productions^19-21^, reflecting a form of rapid, trial-by-trial sensorimotor learning.

**Figure 1.**
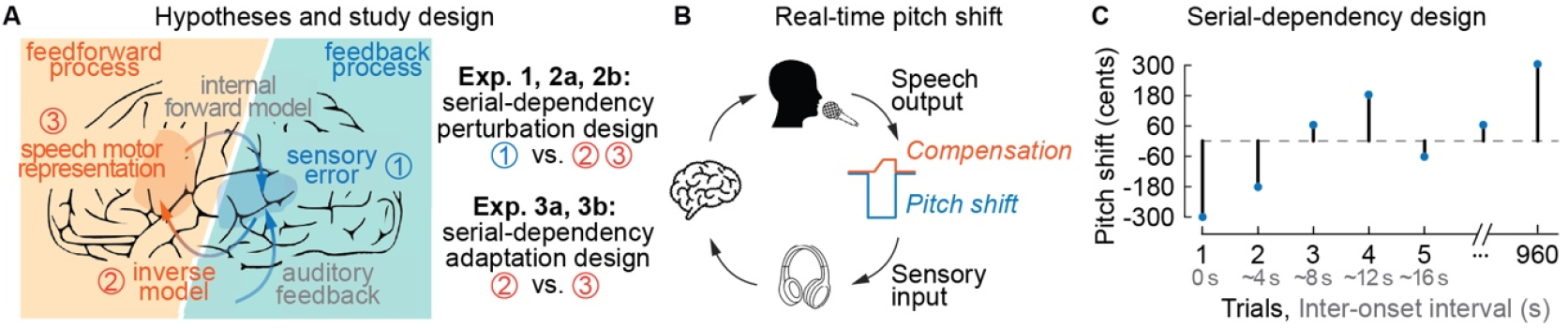
Study design. **A**, Schematics of a processing model of speech production control. Feedback and feedforward processes interactively mediate the control of speech production. When auditory feedback is perturbed and is inconsistent with the predicted consequence from an internal forward model, sensory error is generated and triggers the inverse model in the feedback control. The output of the feedback control further updates the programming of speech-motor representation to compensate for the auditory perturbation. The auditory perturbation and resultant motor compensation could leave traces in either feedback and/or feedforward control processes that mediate sensorimotor learning, even on a trial-by-trial basis. We separately use two paradigms, firstly to test whether sensorimotor learning arises from the changes in the feedback or feedforward processes, and secondly to test whether the learning can be attributed to the updates in the inverse model or in the speech motor representation. **B**, Real-time pitch perturbation during vowel production. The pitch of auditory feedback is artificially shifted (blue) from the utterance recorded in the microphone (orange). The shifted auditory feedback is sent to earphones, which causes compensation in the opposite direction of the pitch shift (the upward change in the orange line of the utterance after the downward pitch shift in the blue line). **C**, The trial sequence of the experiment. Participants undergo 320 trials each day for three consecutive days. The perturbation amounts and directions are randomly applied to each trial.

It remains heavily debated, however, how these serial adjustments contribute to sensorimotor learning via feedback and feedforward control mechanisms.^5,22-25^ One possibility is that serial adjustments reflect changes in the feedback control process, whereby sensory errors encountered in a previous utterance recalibrate the sensory system’s sensitivity to subsequent errors.^26,27^ Supporting this hypothesis, recent findings show that exposure to unreliable auditory feedback down-weights auditory error signals and leads to reduced compensatory response in subsequent production.^19,28^ Alternatively, another possibility posits that serial learning reflects updates to the feedforward process, and motor compensation acts not only as an immediate corrective response, but also leaves short-term traces in the speech sound map or articulator map (as proposed in the DIVA model).^22,25,29,30^ These traces can persist beyond the removal of perturbation, leading to relatively long-lasting adaptive changes that shape subsequent productions even in the absence of feedback perturbation and sensory errors.^20^

Yet, disentangling the contribution of feedback and feedforward processes to sensorimotor learning remains a major challenge, as sensory errors and motor compensations are tightly coupled. Standard auditory feedback perturbation and adaptation paradigms struggle to differentiate the two because of their consistent co-occurrence across trials.^31^ To overcome these obstacles, we employed a serial-dependence paradigm that leveraged trial-by-trial variability to dissociate the effects of past sensory error and motor compensation on future speech behavior, allowing us to test feedback and feedforward hypotheses (**Fig. 1A**, right panel).^32,33^ In Experiments 1 and 2 with a serial-dependency perturbation design, if previous sensory errors did not influence compensation in the current trial, this would suggest that the ‘one-shot’ serial learning occurs either in the subsequent inverse model or within the speech motor representation. In Experiment 3 with a serial-dependency adaptation design, perturbation was introduced in the previous trial but not in the current trial; the inverse model would not operate due to the absence of perturbation. If behavioral changes were still observed despite the absence of perturbation, this would indicate that serial learning effects are due to updates in the speech motor representation within the feedforward process. Our results in five experiments consistently show that prior motor compensation, rather than sensory error, predicts subsequent vocal production. The serial learning effects from past perturbed production adapt to the current production without perturbation. These results provide evidence indicating that motor compensation is not only a reflexive response to sensory perturbation, but also drives updates in speech motor representation, forming an instantaneous plasticity under the linguistic constraints during continuous speaking.

## Results

### Experiment 1: Trial-to-trial learning in repeated production of identical vowel

In the first experiment, we used the simplest setting where participants repeatedly produced a single vowel continuously for 2.5 sec (*N* = 14). When their auditory feedback (**Fig. 1B**, blue lines) was unexpectedly downward or upward perturbed, subjects could compensate for the shift in speech output (orange lines). Through a long sequence of trials in which six levels of perturbation amount were randomly applied to each trial (**Fig. 1C**), we separately measured how the serial changes of the current trial depended on the sensory error and motor compensation of the immediately preceding trial. Throughout the study, we defined each within-trial perturbation and compensation as C_t_ and P_t_, and its preceding-1-trial perturbation and compensation as C_t-1_ and P_t-1_.

We first measured motor responses to perturbation in the current trial (i.e., the relationship between C_t_ and P_t_). We averaged across trials with the same amount of perturbation. It demonstrated that motor compensations were opposed to the direction of pitch perturbation, started around 150 ms after the onset of the perturbation in the conditions of -300 cents, -180 cents, -60 cents, and 60 cents pitch shift (**Fig. 2A**, from left to right, respectively, *ts*(13) = 36.13, 56.36, 109.85, and -92.86, *ps* = 0.043, 0.021, 0.008, and 0.024, cluster-based permutation test), yet this effect was not significant in the condition of 180-cent and 300-cent perturbations (*ps* > 0.203). To compare conditions, we selected a 150∼250 ms window that was commonly used to quantify the amount of motor compensation.^21,34^ Compensation amounts did not significantly differ across upward and downward perturbations (*p* > 0.067, Friedman’s chi-square test, FDR corrected; **Fig. 2B**).

**Figure 2.**
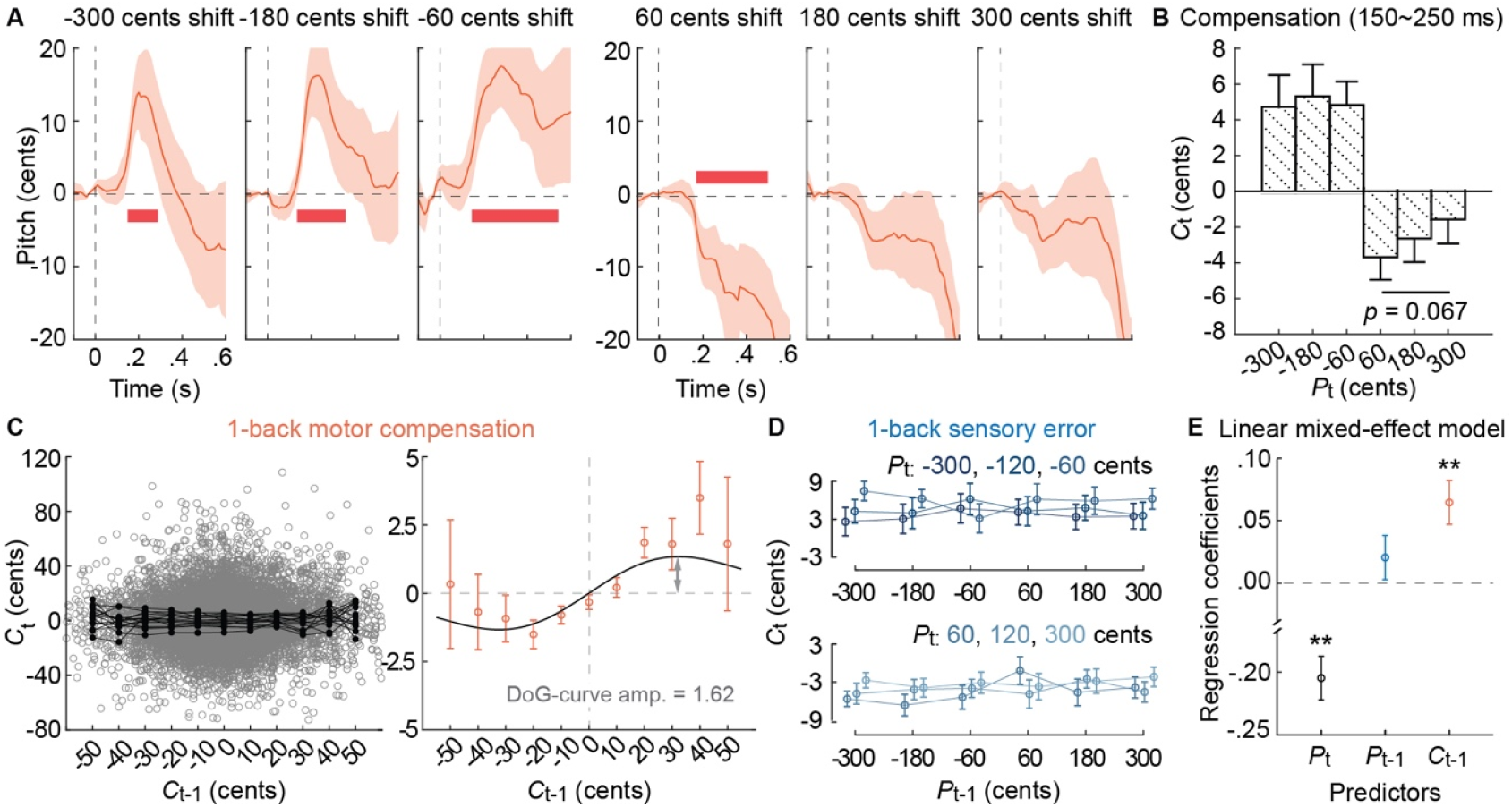
Serial changes of the immediately preceding trial on the compensatory responses of the current trial. **A**, Pitch traces of motor compensation in response to upward or downward perturbations. The orange lines denote the averaged pitch trace across participants. The red horizontal bars denote the times of the significant effects of compensation (*p* < 0.05). **B**, Temporal averaged compensation in the periods of interest (150-250 ms after the perturbation onset). In **C** and **D**, we plot compensation of the current trial, C_t_, as a function of compensation in the immediately preceding trial, C_t-1_, and as a function of the perturbation in the immediately preceding trial, P_t-1_, respectively. **C**, In the left plots, each dot represents one trial and each black curve represents one participant. The right plots show the relationship between preceding and current trials after averaging across trials and participants. Color dots projecting to the same sign on the x and y axes indicate that C_t_ is in the same direction as C_t-1_. **D**, C_t_ as a function of P_t-1_ does not show significant effects, suggesting the current compensation is not influenced by the perturbation amount of the immediately preceding trial. **E**, The effects of current perturbation (P_t_), preceding perturbation (P_t-1_) and preceding compensation (C_t-1_) on current compensation (C_t_) revealed by a generalized linear mixed-effect model. Regression coefficients for each predictor are extracted from the model. Current perturbation and preceding compensation exert opposite and attractive influences, respectively, on the current compensation. Error bars denote SEM across trials. In **A**-**D**, shaded areas and error bars denote the standard error of mean (SEM) across participants. \**p* < 0.05, \*\**p* < 0.005.

Next, we tested the feedforward and feedback hypotheses by separately measuring the dependency of preceding-1-trial sensory error or motor compensation on its subsequent trial. Specifically, we characterized whether C_t_ (in each trial) would follow or oppose C_t-1_ or P_t-1_ (in its preceding trial). Therefore, we plotted C_t_ as a function of C_t-1_ and P_t-1_, respectively (**Fig. 2CD**). The averaged data across trials (grey dots) and participants (black lines) demonstrated that functions of C_t_ to C_t-1_ yielded a derivative-of-Gaussian-shaped (DoG) curve (ps = 0.0001 for both, two-sided bootstrap, FDR corrected) (**Fig. 2C**). That is, C_t_ in the current trial followed C_t-1_ in the preceding trial, especially around 0; if C_t-1_ were approaching the extreme condition (far from 0), the following effect became weak.

We further combined trials with the same P_t_ to isolate the dependency of P_t-1_ (in the preceding trial) on C_t_ (in the current trial). The main effect of P_t-1_ was not significant (χ^*2*^(5) < 6.65, *p* > 0.247; Friedman’s chi-square test, FDR corrected). We further built a linear mixed-effect model to validate the contribution of P_t_, P_t-1_, and C_t-1_ to the C_t_. Consistently, the results revealed a significantly negative coefficient of P_t_ (same as **Fig. 2A**) and a significantly positive coefficient of C_t-1_ (same as **Fig. 2C**) (*t*(11,680) = -21.8 and 3.03, p = 0.0001 and 0.004, respectively, one-sample *t*-test, FDR corrected; **Fig. 2E**). No effect was found for P_t-1_ (same as **Fig. 2D**) (*t*(11,680) = 1.55, *p* = 0.160; **Fig. 2E**). These results suggested that the preceding sensory error only contributed to the instant computation for compensation, but not directly to the future production. It also suggested that updates in the feedforward control process, not the feedback process, regulated subsequent production.

### Experiment 2: Trial-to-trial learning across the production of different vowels

Speech production involved variable vowel categories and thus in Experiment 2, we explored whether the serial effect of C_t-1_ remained across different vowels. In Experiment 2a, participants (*N* = 17) were asked to produce four different vowels that were randomly presented in each trial, i.e., 啊 (/a1/), 咿 (/i1/), 呃 (/e1/), and 哦 (/o1/). We observed a similar serial effect of C_t-1_ on C_t_ (*p* = 0.0001, two-sided bootstrap, FDR corrected; **Fig. 3A**), but not for the effect of P_t-1_ (**Fig. S1**). When comparing experiments 2a and 1, the amplitude of the DoG curve was significantly lower when participants produced different vowels than when they produced the same vowel (p = 0.036, two-sided bootstrap, FDR corrected; **Fig. 3C**). The reduction of serial learning can be caused by different vowels or by the uncertainty of next vowel. To exclude the latter possibility, we conducted Experiment 2b where the four vowels were presented in a fixed order; that is, participants would anticipate the identity of the next vowel. We found a consistent serial effect caused by C_t-1_ (*N* = 12; *p* = 0.0001, two-sided bootstrap, FDR corrected; **Fig. 3B**). Importantly, the amplitude of the DoG curve during fixed-order vowel production was similar to that during random-order (*p* = 0.307, two-sided bootstrap, FDR corrected; **Fig. 3C**), suggesting that serial learning of motor compensation was reduced due to different vowels, instead of uncertainty of next production.

**Figure 3.**
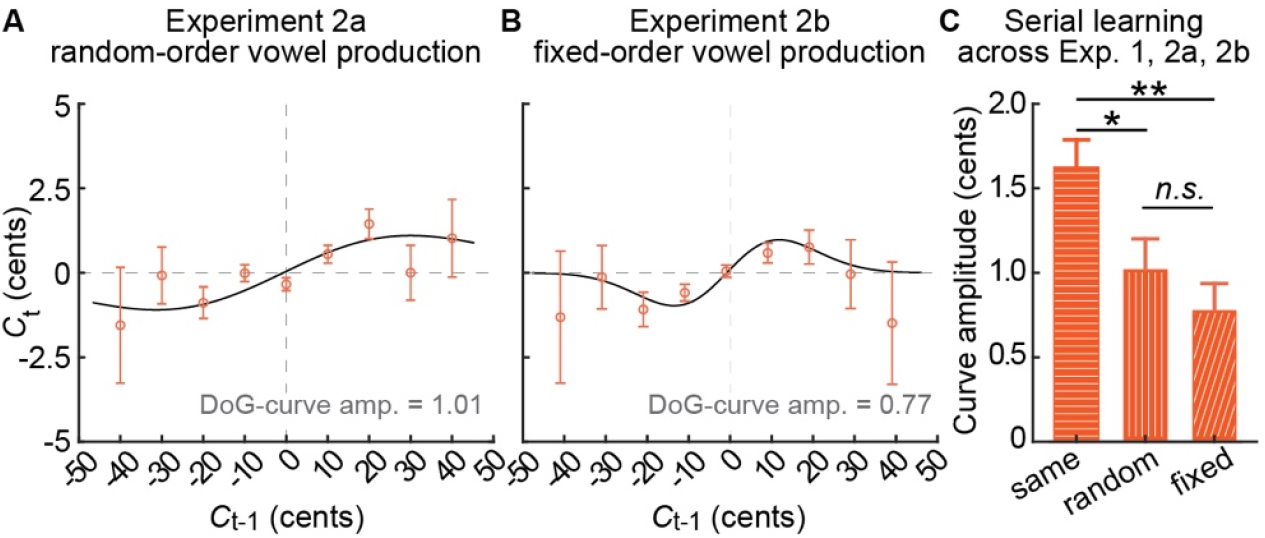
Serial effects across different vowel categories. **A** and **B**, C_t_ as a function of C_t-1_ when participants produce randomly-presented vowels (**A**) and when participants produce expectedly-presented vowels (**B**). In both experiments, significant serial changes to the current motor compensation are observed. **C**, The amount of compensation, characterized by the amplitude of the DoG curve, shows differences between experiments 1, 2a, and 2b, where participants produce the same vowel, different vowels with uncertainty, and different vowels with certainty, respectively. The learning effects are weaker when producing different vowels, compared with those of the same vowels. Error bars denote one SD of the bootstrap distribution. \**p* < 0.05, \*\**p* < 0.005. See also Figure S1.

### Experiment 3: inter-syllabic learning in sentence production within and across word boundary

Experiments 1 and 2 demonstrated that previous motor compensations drove trial-to-trial updates in current speech production, indicating that experience influenced future productions via the feedforward, rather than feedback, control process. However, it remains unclear whether this compensation-driven learning reflects updates in the inverse model or direct modifications to the speech motor representation itself (**Fig. 1A**). To resolve this ambiguity, we employed an adaptation paradigm in which auditory perturbation was selectively applied to an initial syllable, while leaving the subsequent syllable unperturbed. If trial-to-trial learning relies on updating the inverse model to correct sensory errors, then no adaptive changes should emerge during the unperturbed syllable since the inverse model would not be activated without sensory error.

Alternatively, if previous compensations directly reshape the speech motor representation, adaptive changes should persist in subsequent unperturbed syllables regardless of sensory error.

We manipulated the auditory feedback during the continuous production of sentences at different positions of syntactic structures in two separate experiments (*N* = 15 and 13 in Experiment 3a and 3b, respectively). During sentence production, auditory pitch perturbation was randomly applied to either the second, third, or fourth syllable in each trial (**Fig. 4AC**). We quantified compensation in the perturbed syllable and its serial changes in the next syllable, which could be within or across the word boundaries (shown as grey rectangles in insets; **Fig. 4BD**). We showed that upward pitch perturbation drove a significant compensatory response around 200 ms after perturbation onset, compared with the pitch trace without perturbation (*t*(14) < -40.30, *p* < 0.009, cluster-based permutation test; **Fig. S2**). We extracted 200-300 ms mean pitch as compensation amount in the perturbed syllable and as serial-learning amount in its next syllable. Only when the two syllables formed a word, the compensation in the preceding syllable influenced the production of the current syllable (right plot in **Fig. 4B:** *r* = 0.09, *p* = 0.003; left plot in **Fig. 4D**: *r* = 0.22, *p* = 1 × 10^−11^; Spearman correlation).

**Figure 4.**
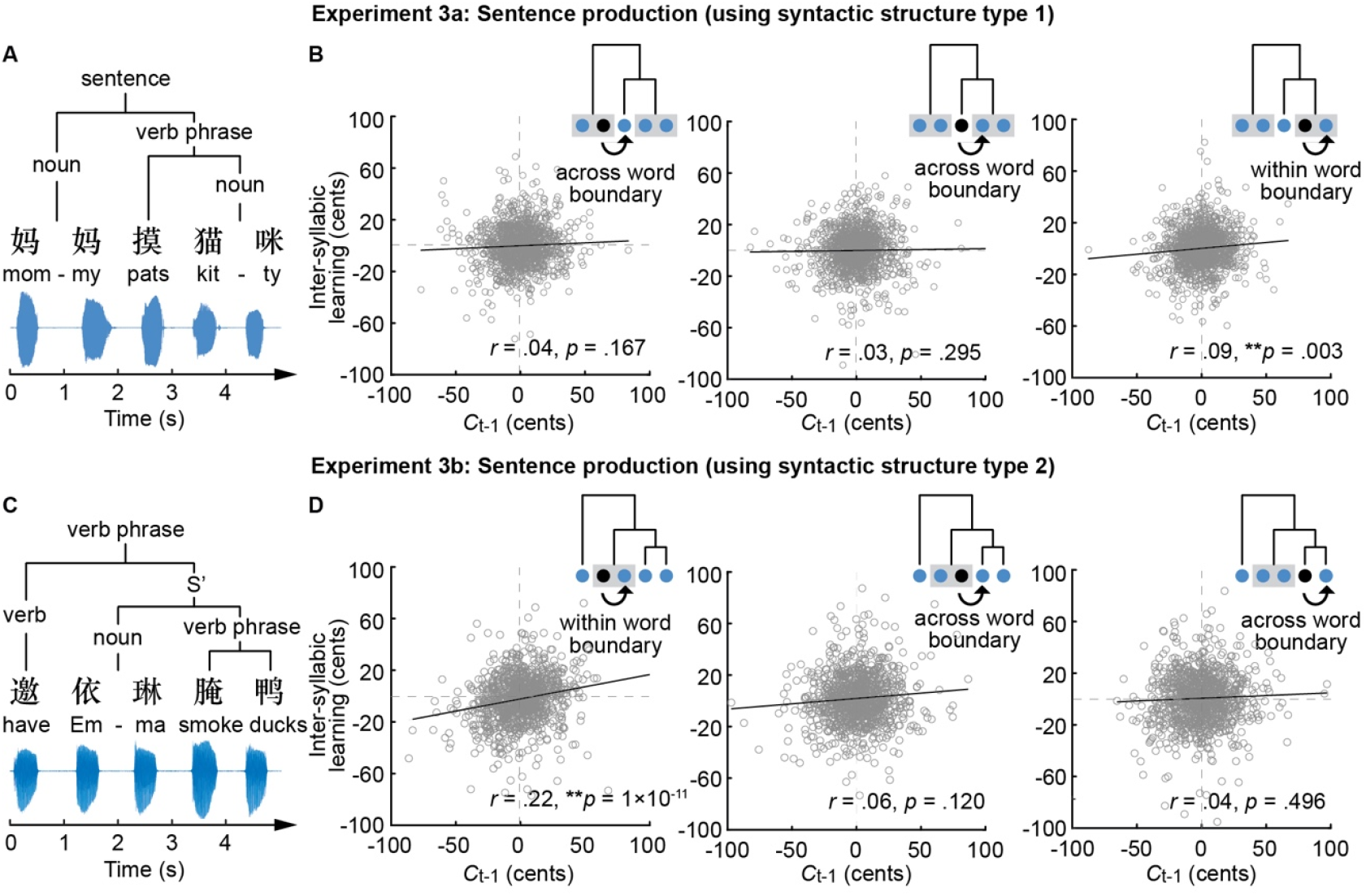
Inter-syllabic learning persists only within the word boundary. **A** and **C**, Experiment 3a and 3b. Participants are asked to produce three 5-syllable sentences that have the same syntactic structure in each experiment. The position of the two-syllable noun is either at the end (Exp. 3a) or in the middle of the sentence (Exp. 3b). Each character is presented on the screen for 500 ms, followed by a blank for 500 ms. **B** and **D**, Relations between the compensation amount of the preceding syllable (x-axis) and serial-learning amount of the current syllable (y-axis) in Exp. 3a and 3b. The mean pitch averaged across 200-300 ms is separately extracted from the perturbed syllable (black dots in the insert plots) and the subsequent syllable (the dot pointed by a black arrow). The grey-shaped rectangle denotes the two-syllabic word. Only when the two syllables are within a word boundary, significant inter-syllabic learning is observed. **p < 0.005. See also Figures S2 and S3.

The correlation coefficients were larger in the within-word-boundary condition than in the across-word-boundary condition (**Fig. 4B**: *p* = 0.180 and 0.026 for position 2 vs 4 and for position 3 vs 4, respectively; **Fig. 4D**: *p* = 0.010 and 0.0005 for position 2 vs 3 and for position 2 vs 4, respectively; one-sided bootstrap, FDR corrected). In contrast, auditory perturbation in the preceding syllable did not contribute to inter-syllabic learning (**Fig. S3**). These results strongly suggested that compensation-driven learning persisted in the current vocal production even without any perturbation and compensation, and the serial learning was constrained by word boundary. It also suggested that past compensations directly modified speech motor representations for the instantaneous, trial-to-trial learning.

## Discussion

We investigated how previous sensory error and motor compensation regulated trial-to-trial learning. We showed that motor compensation in the preceding trial influenced speech production in the next trial, whereas sensory errors did not have any effects.

Crucially, this compensation-driven serial learning was modulated by linguistic contexts -- persisting across different vowels in a string of production and across vowels embedded within the word boundary. These results suggested that instantaneous sensorimotor learning was driven by motor compensation and was constrained by the linguistic system in continuous speech.

The study introduces a serial-dependence paradigm in the domain of speech production that dissociates the influence of previous sensory error and motor compensation on the current vocal production. We observed that when the compensation in trial t-1 was around zero -- despite the presence of sensory error -- the compensation in trial t also returned to baseline. Whereas, when the compensation in trial t-1 was positive or negative, compensation in trial t followed the same direction. These observations are consistent in all three experiments (**Figs. 2C, 3AB**), and can even be found in individual subjects. Moreover, the compensation-driven serial learning was also observed in the production of syllables without feedback perturbation (**Fig. 4**). This evidence suggests that instantaneous plasticity in speech production emerges primarily through immediate updates in the feedforward control process, initiated by preceding compensatory responses. Using a novel serial dependency paradigm, this study extended sensorimotor learning in a relatively longer time scale^32,35,36^ to trial-by-trial instantaneous plasticity.

The distinction between sensory error and motor correction on driving adaptation has been discussed in the motor control literature. For example, studies on reaching have shown that behavioral adaptation occurs even when online motor corrections are unavailable, suggesting that sensory errors suffice to update subsequent movements.^37^ Speaking, however, differs from limb movements in many aspects, including the automatic and involuntary nature of sensorimotor mappings,^38,39^ as well as its mandatory interplay with higher-level linguistic processing. It is possible that speech corrective signals are generated in response to sensory errors in the brain and drive adaptive changes in subsequent productions, regardless of overt compensatory behavior.

Moreover, the storage-based hierarchical yet fixed mappings between the linguistic and motor systems may reinforce the feedforward process during control and learning in speaking, as demonstrated in the results of Exp. 3. Rather than removing compensatory behavior entirely, the current study directly tests whether the amount of compensation in one trial influences the next, providing new insight into the trial-by-trial dynamics of sensorimotor learning in speech.

The relationship between within-trial compensation and next-trial learning is not linear; it is more likely a DoG-shaped curve. Specifically, if the previous compensation is large, the serial changes in the current trial become smaller, suggesting a dynamic adjustment process during speech planning.^5,6,40^ Furthermore, this process is also constrained by the similarity of production targets between trials. If the target in the next trial changes to a different vowel, the trial-to-trial learning becomes weaker (**Fig. 3C**). Rochet-Capellan et al. ^41^ tested whether sensorimotor learning during training transfers to new utterances with different sounds and demonstrated that it is highly dependent on the specific utterances. That is, the experience in the motor domain becomes less useful when the next utterance is different.

Lastly, the extension of compensation-driven serial learning into sentence production reveals that serial learning only exists within word boundary (**Fig. 4**). In Experiment 3, participants may simply read out aloud each visually presented word without syntactic processing. If this were true, inter-syllabic learning, as observed in Experiments 1 and 2, would occur between every two syllables. However, the observed significant effects of syntactic structures are inconsistent with the alternative ‘reading-only’ hypothesis.

Moreover, the extensive training on the three 5-syllabic sentences ensured that participants were extremely familiar with the materials and retrieved the simple sentential structure at the visual presentation of the first word. All the experimental settings and positive results support the hypothesis regarding the role of syntactic constraints in compensation-driven serial learning.

The results of within-word-boundary inter-syllabic learning suggest that words serve as a central unit in speech planning. Motor compensation in the preceding syllable can be updated to the next within-word syllable. However, any compensation-based experience is not available for the syllables across words. Psycholinguistic studies, such as those by Levelt ^42^, propose that during speech production, words are retrieved from the mental lexicon as a whole unit, followed by phonological encoding into syllabic and phonetic representations. The entire course of speech production at the linguistic level would interact with sensorimotor processes via internal forward models and regulate feedforward processes at both featural and temporal domains,^11,13,43-47^ to achieve efficient online control. The observed serial effects within word structure suggest that the programming and regulation of speech production are presumably in the size of the encoding unit within the word structure, which is consistent with the hypothesis of interaction between linguistic and sensorimotor systems in speech production control.^4^

The study showed that previous motor compensation can modulate the subsequent vocal pitch production; whether this compensation-driven serial learning can be generalized to other speech domains, such as formants^19^ and intensity^11^ that are controlled by distinct vocal apparatus and may be mediated by potentially different mechanisms, requires further investigation. Considering a broader linguistic context in speech motor control is challenging and relevant studies are rare.^48,49^ In this study, we used pitch as a representative dimension and carefully selected different combinations of consonants and vowels in continuous speech to pioneer the investigation. Future studies should be carried out with caution and careful controls. For example, pitch contours in continuous speech can be potentially interfered with by consonantal perturbations^50^ or sentence intonation^51^. During the production of non-sonorant consonants, the vibration of vocal folds will be affected, leading to a break and a carry-over rising or falling in the pitch contours.^52^

In summary, by implementing a novel serial-dependence paradigm in the domain of speaking, we found distinct functions of sensory error and motor compensation in sensorimotor learning in continuous speech. Motor compensation drives updates in the feedforward control process and regulates trial-to-trial learning that was constrained by linguistic contexts, while sensory error only contributed to the instant computation of compensation. These results suggest that the history of motor processes significantly contributes to sensorimotor learning and control. Systematic interplay among cognitive, motor, and sensory systems regulates behaviors.

## Materials and methods

### Participant

In five experiments, i.e., Experiments 1, 2a, 2b, 3a, and 3b, we recruited 14 participants (9 females, 18-27 years old, mean 21 years old), 17 participants (11 females, 19-32 years old, mean 23 years old), 12 participants (8 females, 20-27 years old, mean 22 years old), 15 participants (9 females, 19-25 years old, mean 22 years old), and 13 participants (7 females, 18-26 years old, mean 21 years old), respectively. They were college students with self-reported normal hearing and speech. All participants were right-handed and were native Chinese speakers. Each volunteer only participated in one experiment. Written informed consent was obtained from each participant before the experiment. The experimental procedures were approved by the Research Ethics Committee at the East China Normal University and New York University Shanghai.

### Procedure in Experiment 1

Participants were asked to produce the vowel /a/ in a total of 960 trials over three days. Each trial began with a white fixation cross, displayed at the center of the screen for a randomized duration of 0.8–1.2 s (uniform distribution), signaling participants to be prepared for vocalization. When the cross turned green, participants were instructed to vocalize no more than 500 ms and sustain the /a/ sound with a steady and natural tone for 2.5 s until the cross disappeared. A pitch perturbation for a duration of 500 ms was randomly introduced 0.8–1.2 s after vocal onset. The direction (i.e., upward or downward) and magnitude (i.e., 60, 180, or 300 cents) of the pitch shift were randomly assigned across the 960 trials for each participant. At the end of each trial, participants received visual feedback: “Good,” “Too slow,” or “Too short.” Participants completed 95.86% ± 0.02% (s.d.) of trials that met criteria. Each trial was followed by a silent interval with a random duration of 1–2 s (uniform distribution) before the next trial began.

Participants completed 320 trials in 10 blocks each day, with self-paced rest periods of at least 1 minute between blocks. At the beginning of each session, participants underwent a training block of 10 trials. The mean pitch from these trials was used to define the pitch range in the Audapter toolbox (see *Auditory feedback perturbation* below).

### Procedure in Experiment 2a

The procedure for Experiment 2a was identical to Experiment 1, except participants were required to produce four different vowels without semantic meaning—啊 (/a1/), 咿(/i1/), 呃 (/e1/), and 哦(/o1/)—randomly assigned in each trial. A white letter corresponding to the syllable was presented at the center of the screen, and upon turning green, participants were instructed to vocalize a vowel. Participants completed 94.82% ± 0.11% of trials that met the criteria.

### Procedure in Experiment 2b

The procedure for Experiment 2b was identical to Experiment 2a, except the order of four vowels were presented in a fixed order, i.e., /i1/-/a1/-/e1/-/o1/ in every four trials so that participants would not anticipate the next vowel production. Participants completed 97.55% ± 0.03 of trials that met the criteria.

### Procedure in Experiment 3a

The procedure for Experiment 3a followed that of Experiment 1, with two differences. First, participants completed 300 trials in a single day. Second, they were required to produce sentences with the same syntactic structure rather than isolated vowels. In each trial, one of three five-syllable sentences (i.e., ▪▪▪猫咪 [ma1 ma1 lu1 mao1 mi1], 丫邀巫医 [ya1 ya1 yao1 wu1 yi1], 依依摸▪▪ [yi1 yi1 mo1 wu1 ya1]) was randomly selected (uniform distribution), with each syllable appearing sequentially at the center of the screen for 500 ms, followed by a 500 ms blank screen. The whole duration for a sentence production was 5 s. During vocalization, a whole-utterance pitch perturbation (upward or downward perturbation at 60, 180, 300 cents, randomly assigned) was applied to the second, third, or fourth word during vocalization (uniformed distribution). In 20% of the sentences, no perturbation was applied. The speech materials and task were carefully chosen to avoid the effect of consonantal obstruction^53^ and the effect of articulators’ dynamics on pitch contour, respectively. Participants completed 94.44% ± 0.07% of trials that met the criteria.

Before the main experiment, participants completed a training session in which they produced each of the three sentences four times. They were explicitly instructed on the content and syntactic structure of the sentences. Additionally, they were instructed to vocalize each syllable within 500 ms of its appearance and to avoid coarticulation between words. Participants could repeat the training session upon request.

### Procedure in Experiment 3b

The procedure for Experiment 3b was the same as Experiment 3a, except that the three sentences were replaced with another three with a different syntactic structure, i.e., 邀依琳摸 ▪ [yao1 yi1 yi1 mo1 ying1], 拉▪▪腌▪ [la1 ma1 ma1 yan1 ya1], and ▪丫丫溜猫 [yue1 ya1 ya1 liu1 mao1], where the two-syllable noun was in the position of second and third character in the sentences. Participants completed 97.36% ± 0.03% of trials that met the criteria.

### Auditory feedback perturbation

Participants’ speech was recorded by a Shure beta 58A microphone that was placed 5 centimeters away from their mouths. The acoustic signals from the microphone were pre-amplified and then provided to an audio interface (MOTU Microbook IIc), collected at a 48k Hz sampling rate. The recorded speech was real-time manipulated (i.e., perturbation) in the audio interface, and the processed signal was split and sent both to the computer and to the headphones. The speech processing in the audio interface was performed using the Audapter toolbox.^54^ The pitch of speech in each trial was shifted using the time-domain method. A 10-ms linear ramp was added at boundaries to attenuate possible abrupt jumps in the speech. A priori pitch range was set as +/-50 Hz relative to the mean pitch of a vocal /a/ for each participant. The rest of the parameters were set as default including algorithm, frame length, and cepstral liftering.

### Acoustic analysis

In all experiments, we analyzed signalIn, the speech output from participants. In experiments 1 and 2, we first removed silence from the recordings of produced vowels in each trial using the *vad* function. Visual inspection was performed to make sure noise did not affect the voice activity detection. Pitch tracks (at a 100-Hz sampling rate) were measured using the Praat-based script ProsodyPro.^55^ Default parameters were used. Perturbation onset in the pitch trace was obtained by a customized automatic algorithm combined with manual inspection. To convert hertz to cent scale, we defined baseline pitch by taking the mean of pitch trace in the -400 to -200 ms before perturbation onset. The pitch trace was transformed according to the formula: 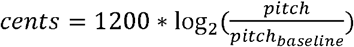. A trial rejection was performed if the pitch trace was varied beyond its perturbation amount minus 100 cents (for downward-perturbation trials) or plus 100 cents (for upward-perturbation trials). To extract perturbation and compensation amounts for DoG-curve fitting and linear mixed effect model, the pitch trace in the 50-80 ms and the 150-250 ms windows were averaged, respectively, for each trial.^21^ Compensation amounts for all trials and all participants were corrected by subtracting the grand average. This correction ensures that the DoG-curve fitting starts from zero, allowing its amplitude to be accurately measured and compared across experiments.

In Experiment 3, we applied the same pipeline except that only the perturbed syllable and its subsequent syllable were extracted to measure their pitch trace. Based on the visual inspection of pitch contours (**Fig. S2**), Mandarin tone 1 produced in isolation was characterized by a high-level tone and showed a pitch contour that typically involved a slight rising intonation at the onset, lasting around 0-150 ms, before stabilizing into a steady high pitch.^56^ Therefore, we calculated the compensation amount in the perturbed syllable and serial-learning amount in the following syllable using mean pitch in the 200-250 ms time window, with baseline pitch in the 150-200 ms time window.

### Statistical analysis of compensatory responses

For participant-level analysis, trials with the same perturbation condition were averaged, regardless of vowel categories in Experiments 2 and 3. To assess whether there was a compensatory response to perturbation, a two-sided cluster-based permutation test^57^ was used to test averaged pitch traces against perturbation amount or between perturbed and normal conditions (**Fig. 1A** and **S2**). A one-sided one-sample t-test was used to test motor compensation extracted from time of interests against 0 (**Fig. 2B)**.

### Statistical analysis of trial-to-trial learning effect

To explore the serial effect of preceding-1-trial motor compensation amount on the current motor response, C_t_ was expressed as a function of C_t-1_. For positive values of this C_t_ or C_t-1_, participants compensated upward in response to perturbation in the current or preceding-1-trial. Consequently, data points in the scatter plot (**Fig. 2C** and **3AB**) that had x- and y-values in the same sign indicated that C_t_ was in the direction of C_t-1_. We assess the trial-to-trial learning effect by polling the compensation amount of all trials and fitting the DoG-shape curve, a well-found psychophysical function in previous serial-dependence studies.^32,36^ The DoG was given as 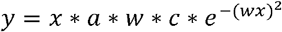, where *a* was the curve amplitude, *w* was the curve width, and *c* was the constant 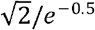. To assess the serial effect of preceding motor compensation, *x* was C_t-1_, and width parameter *w* of the curve was constrained to a range of plausible values (*w* = 0.019-0.071, corresponding to the curve peak between 10 and 36 perturbation pitch difference). The model fitting used the function *fmincon* with default parameters. We employed a bias-corrected and accelerated bootstrap to evaluate the significance of the DoG models and to compare DoG models between conditions. We applied a fitted DoG model to the pooled group data and generated a permutation distribution for the DoG-curve amplitude parameter.^58,59^ This permutation process was repeated 2,000 times. For a one-sided comparison of curve amplitude, if the data population in one condition was greater than A% of the data population in the other condition, the significance level was (100A + 1)/2,001. For a two-sided comparison, the significance level was (200A + 1)/2,001. A false discovery rate (FDR) correction was applied.

To explore the serial effect of sensory error on C_t_, C_t_ was expressed as a function of P_t-1_. We use a Friedman’s chi-square test for multiple condition comparison, i.e., controlling the current perturbation as the same to explore the effect of preceding perturbation on the current compensation (**Fig. 2D**). Spearman correlation was used to find the relation between C_t-1_/P_t-1_ and inter-syllabic learning within and across syntactic boundaries (**Fig. 4BD** and **S3**). To compare the correlation coefficient in within- and across-boundary conditions, a bias-corrected and accelerated bootstrap was used.^60^

### Generalized linear mixed-effect model

To quantitatively test the relations of C_t_ to P_t_, P_t-1_ and C_t-1_, we built a generalized linear mixed-effect models (**Fig. 2E** and **S1**). The model assumed that P_t_, P_t-1_ and C_t-1_ together influenced C_t_. The function ‘fitglme’ was performed and accounted for group-level fixed effects and random effects because of vocal pitch variation across participants. The model was specified as *C*_*t ij*_,= *f* (*β*_0*j*_ + *β*_1*j*_*P*_*t ij*_ + *β*_2*j*_*P*_*t*−1*ij*_ + *β*_3*j*_*C*_*t*−1*ij*_+1|*subjects*), where *i* indicated participant index and *j* indicated *j*^th^ trial. The regression coefficient of each predictor, i.e., *β*, was accessed by the *t*-statistics against the null hypothesis test that the coefficient was equal to 0. FDR correction was applied.

## Data and code availability

All data and code that support the findings of this study is available in Mendeley Data: 10.17632/pybdtwxvzz.2.

## Acknowledgments

We thank Yuan Xie and Bao Hong for their thoughtful comments on previous versions of the manuscript, Tingting Wang and Huixuan Liu for assisting in collecting data. This study is supported by the National Natural Science Foundation of China 32271101, Program of AI-Driven Initiative to Promote Research Paradigm Reform and Empower Disciplinary Advancement by Shanghai Municipal Education Commission (SMEC), Program of Introducing Talents of Discipline to Universities, Base B16018, and NYU Shanghai Boost Fund to X.T.

## Author contributions

Y.L., X.T., and J.C. designed the experiment. Y.L., W.T., and

Z.X. collected the data, Y.L., Z.X., A.X., and W.T., performed the analyses, Y.L., and X.T. drafted, reviewed, and corrected the manuscript. X.T. and J.C. supervised the project.

## Declaration of interests

The authors declare no competing financial interests.

## Declaration of generative AI and AI-assisted technologies in the writing process

During the preparation of this work, the authors used ChatGPT to proofread texts. After using this tool or service, the authors reviewed and edited the content as needed and took full responsibility for the content of the publication.

## Notes

### Competing Interest Statement

The authors have declared no competing interest.

### Summary of Updates

We have significantly revised Introduction and Results.

