## Supplementary figures for "Regulation of sensorimotor serial learning in speech production by motor compensation rather than sensory error"

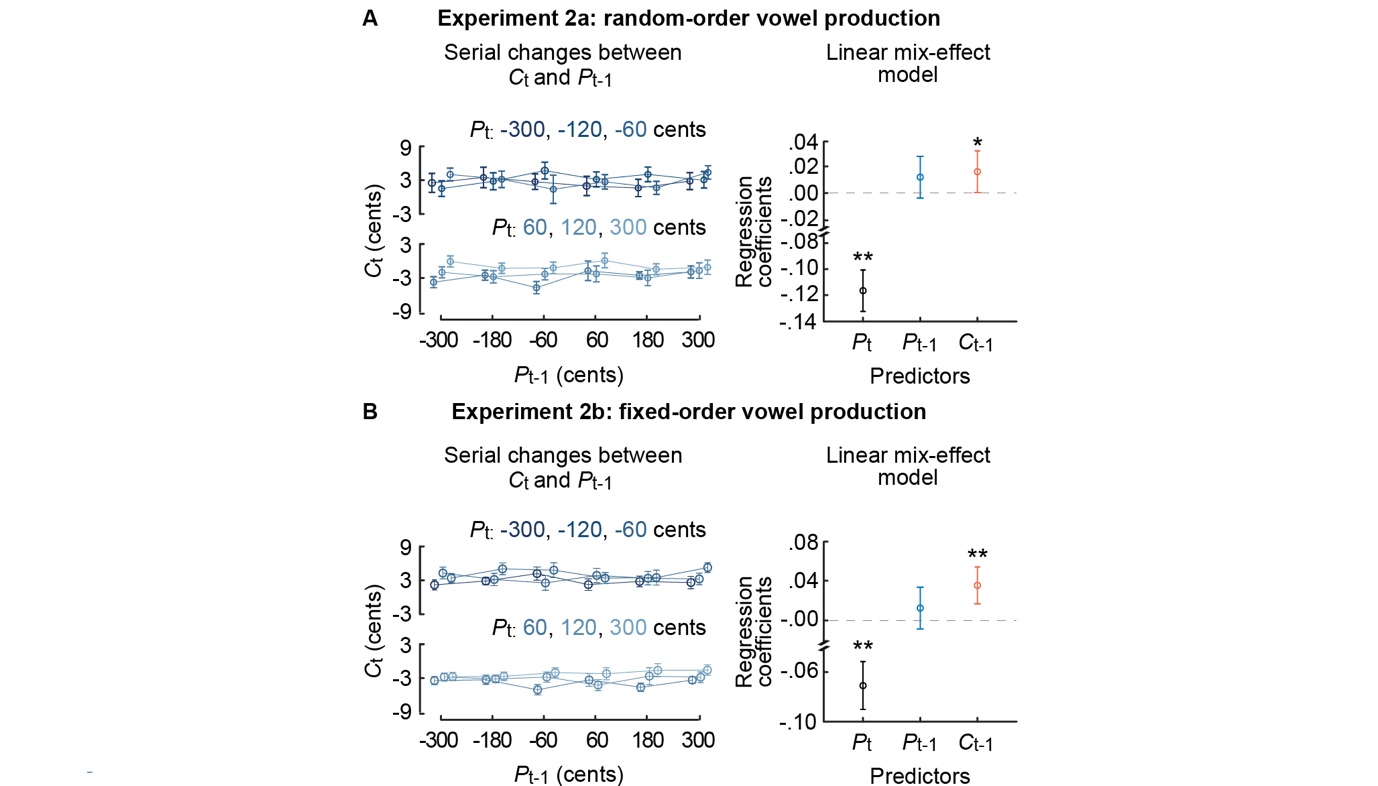

**Figure S1. Sensory error in the preceding trial and compensation in the current trial.** Experiments 2a (**A**) and 2b (**B**) did not obtain any significant relation between P_t-1_ and C_t_. Left: C_t_ as a function of P_t-1_. Upper plot statistics: *χ^2^*(5) < 5.51, *p* > 0.163; Friedman's chi-square test, FDR corrected; Lower plot statistics: *χ^2^*(5) < 6.27, *p* > 0.111; Friedman’s chi-square test, FDR corrected. Right: Regression coefficients of P_t_, P_t-1_ and C_t-1_ on C_t_ are extracted from a generalized linear mixed-effect model. Upper plot statistics: from left to right: *t*(14,994) = -14.36, 1.59 and 2.11, *p* = 0.0001, 0.111, and 0.035, one-sample *t*-test, FDR corrected). Lower plot statistics: from left to right: *t*(10,749) = -7.27, 1.29 and 3.57, *p* = 3 × 10^-13^, 0.198, and 0.0004). Error bars denote SEM across trials. **p* < 0.05, ***p* < 0.005.

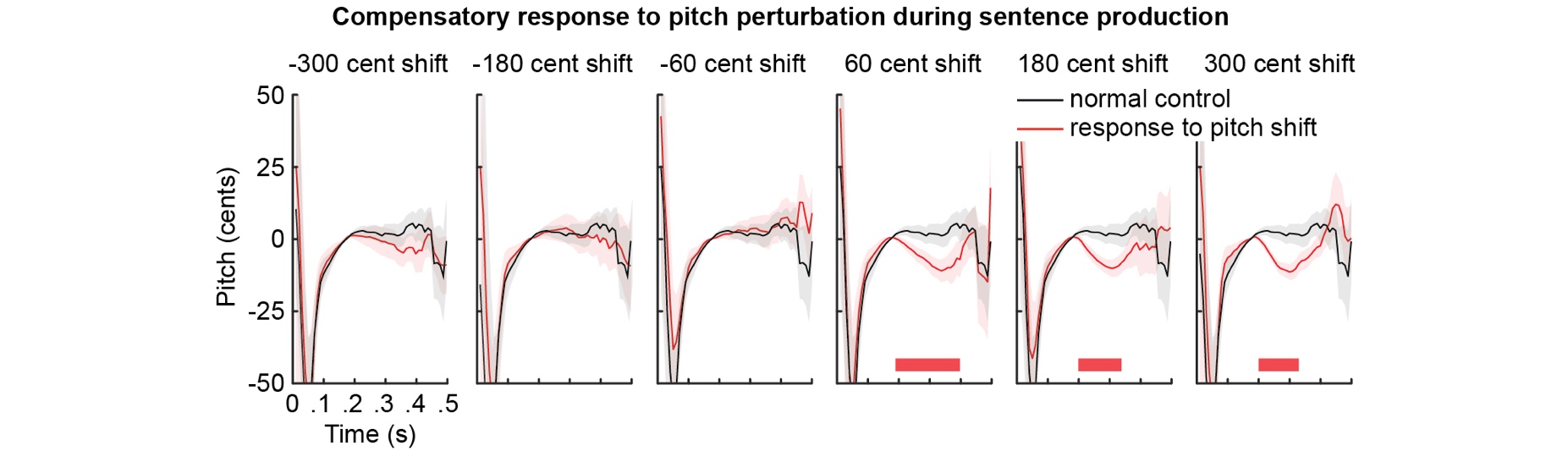

**Figure S2. Motor compensatory response during sentence production.** Pitch traces of perturbed (red curves) and unperturbed words (black curves) averaged across the trials with the same conditions. Grey-shaded areas denote SEM across participants. Red horizontal bars denote the times of the significant effects: 60-cent shift, *t*(14) = -103.52, *p* = 0.0005; 180-cent shift, *t*(14) = -41.05, *p* = 0.007; 300-cent shift, *t*(14) = -40.30, *p* = 0.009, cluster-based permutation test.
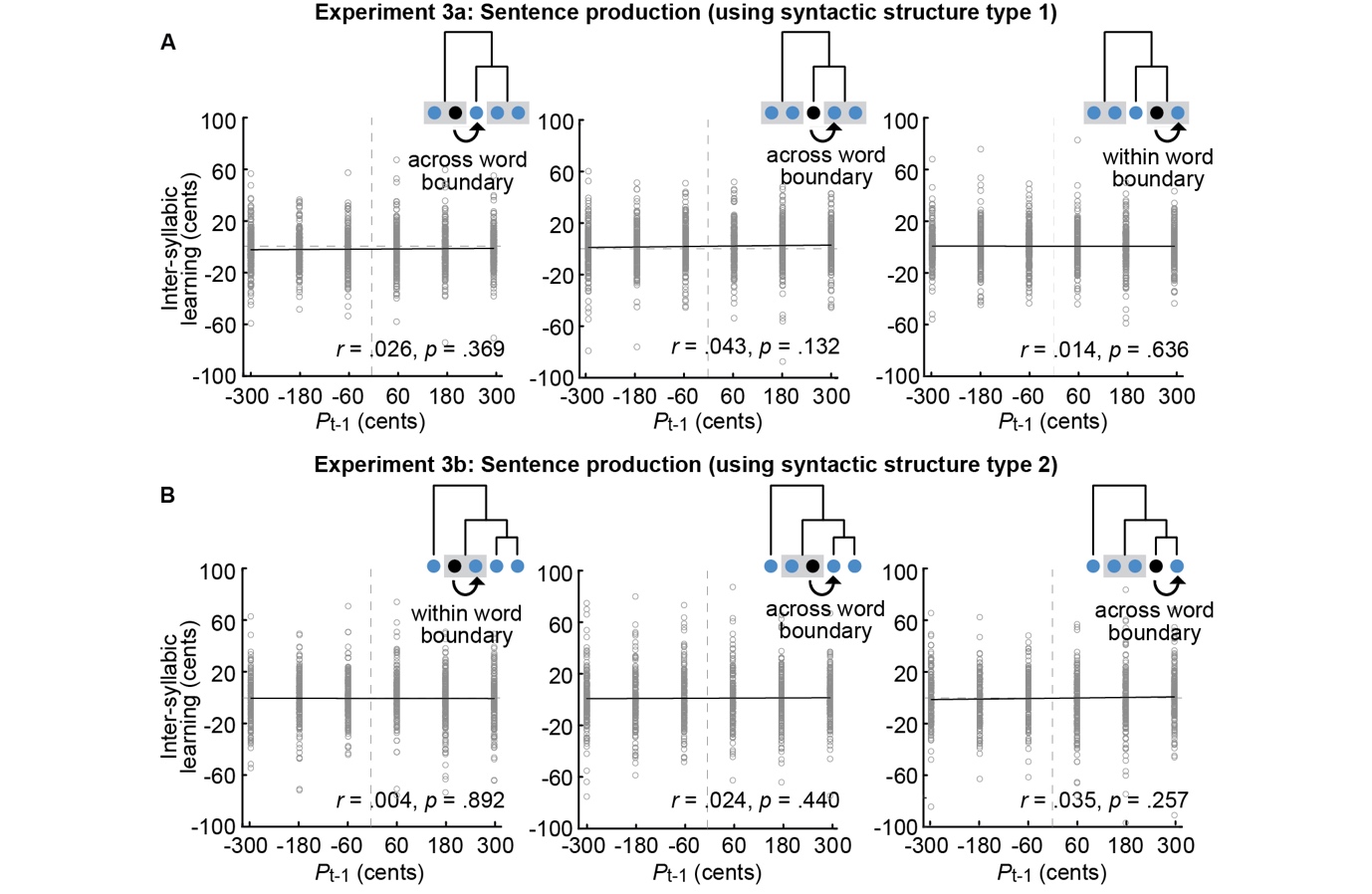

**Figure S3. Inter-syllabic learning is independent of perturbation amount in preceding trial.** In **A** and **B**, we quantify relations between the perturbation amount of the preceding syllable (x-axis) and serial-learning amount of the current syllable (y-axis) in Exp. 3a and 3b. We use experimentally predefined perturbation amount for each perturbed syllable (black dots in the insert plots) and use the same analysis pipeline in Fig. 4 to quantify inter-syllabic learning in the subsequent syllable (the dot pointed by a black arrow). The grey-shaped rectangle denotes the two-syllabic word. For all experiments and syllable positions, auditory perturbation does not contribute to inter-syllabic learning.
